# The blood proteome predicts the impact of circulating factors on age-related mitochondrial health

**DOI:** 10.1101/2025.11.09.687483

**Authors:** Stephanie R. Heimler, Jaclyn Bergstrom, Nina S. Sun, Toshiko Tanaka, Ann Zenobia Moore, Julián Candia, Brian H. Chen, Qu Tian, Arsun Bektas, Giovanna Fantoni, Christopher Dunn, Mary Kaileh, Ranjan Sen, Luigi Ferrucci, Anthony J. A. Molina

## Abstract

Circulating non-cellular factors, such as plasma proteins, contribute to various features of aging. To determine the impacts of endogenous circulating factors on human age-related bioenergetic decline, we treated primary human fibroblasts with serum samples representing the adult life-course. Our results demonstrate that the maximal mitochondrial bioenergetic capacity of fibroblasts treated with serum is negatively correlated with the chronological and epigenetic age of the serum donor. Using targeted proteomics, we identified plasma proteins associated with the bioenergetic effects of serum. We then utilized elastic net, a linear regression modeling technique, to derive a novel proteomic signature of age-related mitochondrial differences. MitoAge is a 25-protein signature of age-related mitochondrial health that predicts the systemic bioenergetic effects of circulating factors and is related to differences in physical function across human aging. Signatures that report on cellular hallmarks of aging, such as mitochondrial function, represent a new generation of mechanistically-informed biomarkers of biological aging.

**GRAPHICAL ABSTRACT:** In this study, we describe the development of a novel proteomic signature, MitoAge. This signature was developed by utilizing human primary fibroblasts treated with serum samples representing the adult human life-course and analyzing how resulting respirometry correlated with chronological age, epigenetic age, and abundance of serum proteins. Utilizing machine learning techniques, we derived a 25-protein signature which can predict bioenergetics and physical features related to aging. Application and utility of this signature may be used to identify novel drivers of health and longevity.

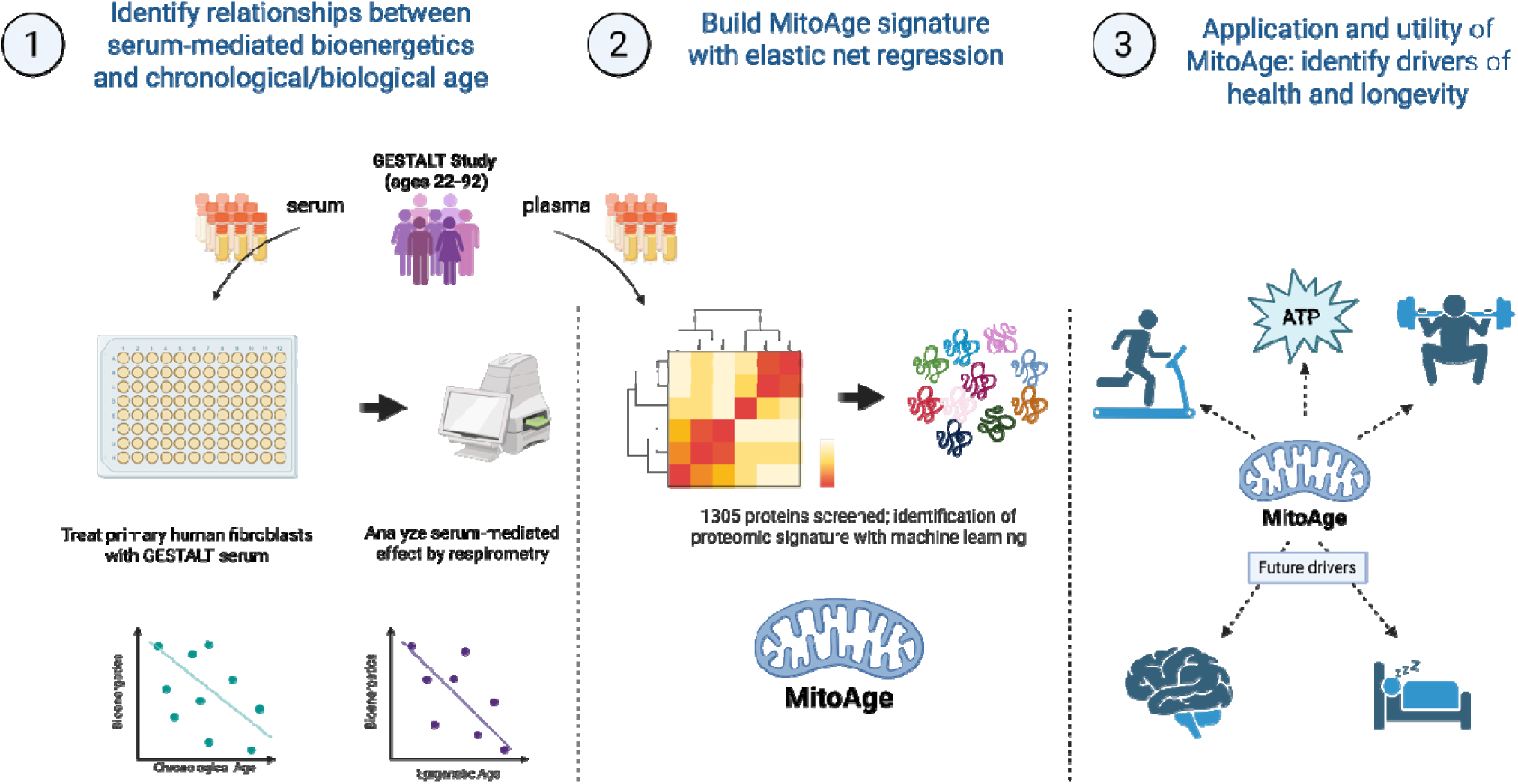

## INTRODUCTION

Aging is one of the strongest predictors of chronic disease, frailty, and a number of adverse conditions that negatively impact older adults. Mitochondrial dysfunction is a widely recognized cellular hallmark of aging (1), and can drive age-related cellular processes (2–5), functional decline (6), and cognitive impairment (7). We and others have shown that age-related mitochondrial differences are systemic, as exhibited by the respirometric profiles of circulating blood cells (8–10).

Multiple lines of evidence demonstrate that blood harbors factors that can drive aging across organ systems, including skeletal muscle (11), liver (12), heart (13), and pancreas (14). Heterochronic parabiosis studies in mice have demonstrated that circulating factors in blood drive multiple features of aging, including mitochondrial dysfunction. Using this model, our group previously reported that blood can mediate skeletal muscle age-related bioenergetic decline (15). Across measures of mitochondrial function, young mice in parabiosis with older mice exhibited mitochondrial deficits associated with aging. Importantly, we observed an equalization of bioenergetic capacities between young and old heterochronic pairs, clearly demonstrating that circulating factors are the primary drivers of age-related bioenergetic differences. We have also reported that circulating factors play a central role in mitochondria changes associated with behavioral interventions, such as diet and exercise (16) and age-related disease, such as Alzheimer’s disease and dementia (17,18). Based on these previous studies, we postulated that in-depth analysis of circulating factors could enable the prediction of systemic age-related bioenergetic decline. Therefore, using plasma from individuals representing the adult human life-course, we sought to identify circulating protein signatures that are predictive of age-associated differences in mitochondrial function.

In recent years, factors found in circulation, such as lipids, metabolites, and proteins, have been associated with chronological age (19–23). Among the types of molecular analytes that can serve as biomarkers, proteins have been shown to be particularly useful, and previous studies have already shown that circulating proteins can serve as promising biomarkers of age and disease (24–28). In order to identify circulating factors that are directly associated with the effects on mitochondrial function, we first treated naïve human primary fibroblast cells with serum from participants representing the adult human lifespan to investigate how mitochondrial oxidative phosphorylation (OXPHOS) correlated with both chronological and epigenetic age of the participants from whom the serum samples were derived. We then conducted proteomic analysis of the serum samples to identify circulating proteins that were most correlated with the bioenergetic effects of these samples. To test the hypothesis that circulating proteins drive the effects of serum on cellular bioenergetics, we utilized computational approaches to develop a signature based on proteomic data alone, and examined how accurately we could predict the bioenergetic effects of whole serum (**Graphical Abstract**). The age-related proteomic signature of mitochondrial health (MitoAge) was trained to predict the bioenergetic effects of serum derived from individuals 22 to 92 yrs of age. Given the recognized role of mitochondria in aging and a wide array of age-related conditions, including physical function decline (29–31), we then examined associations between MitoAge with strength and fitness. The development of signatures that report on cellular hallmarks of aging advances efforts to develop mechanistically informed biomarkers of biological aging. MitoAge was developed to report on age-related bioenergetic differences, relying on the effect of age as part of the model. The goal of MitoAge is to be used across multiple studies to inform on the role of age-related bioenergetic differences and broaden our understanding of drivers of health and longevity.

## METHODS

### 1. Study population

The GESTALT study began in 2015 with the goal of discovering novel biomarkers of human aging. The study enrolled healthy adults aged 20 years and older who were free of major and chronic diseases. Inclusion criteria and participant demographics have been previously described (24). GESTALT protocols were approved by the National Institutes of Health Intramural Research Program Institutional Review Board, and participants provided written informed consent at the time of each visit. The analysis was conducted in 88 participants with serum samples and plasma proteomic data.

### 2. Cell culture and serum treatment

To examine the change in mitochondrial function mediated by serum, we treated two lines of primary human dermal fibroblasts from the San Diego Nathan Shock Center (32,33) with 88 dilute serum samples from GESTALT participants. The two fibroblast lines were from male donors, aged 33 and 72 years old. Fibroblasts were cultured in high glucose DMEM (Thermo Scientific, 11965092), supplemented with 15% FBS, 1% NEAA, and 1% GlutaMAX. For serum treatment, the experimental design was adapted from Gonzalez-Armenta et al. (16). To summarize, cells were seeded in an Agilent Seahorse XFe96 assay plate at 20K cells/well in 160 μL culture media. Following cell seeding, plates were allowed to rest for 45 minutes to 1 hour at room temperature before incubating at 37°C and 5% CO_2_ overnight. The next day, culture media was removed from each well, and 10% GESTALT serum (total in-well volume of 160μL) in culture media was added to each well in quadruplicate. Fibroblasts with serum treatment were incubated for 24hrs before bioenergetic assays. All experiments were conducted between in vitro passage 9-10, and no morphological differences were observed between passages. Cell number per well was quantified using a Gen5 brightfield imager (Agilent, Inc., Santa Clara, CA) pre- and post-treatment, and these quantifications were confirmed by Hoechst nuclear staining. The serum samples used for these exposure assays had corresponding plasma samples, which were used for proteomic analysis (described below).

### 3. Respirometry

To examine the mitochondrial effects of serum treatments, we performed high-throughput respirometry (HTR) on primary human fibroblasts as previously described (16,17). Respirometric analyses were conducted using an Agilent Seahorse XFe96 extracellular flux analyzer (XFe96; Agilent, Inc., Santa Clara, CA). The XF assay media contained XF DMEM Base Medium (Agilent Technologies, 103575-100) supplemented with 1 μM glucose (Agilent Technologies, 103577-100), 1 μM GlutaMAX (Thermo Scientific, 35050061), and 1 μM sodium pyruvate (Thermo Scientific, 11360070). The mitochondrial stress test Seahorse assay was conducted with final in-well concentrations of 1.25 μM oligomycin, 1 μM FCCP, 1.0 μM Antimycin A, 1.0 μM Rotenone, and 32.4 μM Hoechst (nuclear DNA stain; 30-minute incubation). We included control treatments in each plate where cells were cultured either without serum or with 10% FBS. The Seahorse analyzer is calibrated before each assay and is maintained regularly.

### 4. DNA Methylation

Methylation data were gathered from multiple cell types from the GESTALT participants: peripheral blood mononuclear cells (PBMCs), monocytes, and skeletal muscle. The DNA methylation platform used for these analyses was Illumina EPICv1 (human). Blood collection and DNA methylation analysis was conducted as previously described (34). Muscle biopsy preparation was also conducted as previously described (35). Raw DNA methylation data files were processed, annotated, and analyzed using the *SeSAMe* Bioconductor package’s default settings (36,37). Methylation data files were then submitted to the Clock Foundation’s DNA Methylation Age calculator (https://projects.clockfoundation.org/, accessed 2/27/2024), where key outputs were: Original (pan-tissue) Horvath Age (38), DNAmPhenoAge (39), and GrimAge (40).

### 5. Post-exercise phosphocreatine recovery rate (kPCr)

The maximum oxidative capacity of the vastus lateralis (VL) muscle was measured by *in vivo* phosphorus magnetic resonance spectroscopy (^31^P-MRS) and analyzed as previously described (41–43). Briefly, skeletal muscle oxidative capacity was quantified via ^31^P-MRS as post-exercise recovery rate of phosphocreatine (kPCr), which was calculated as the inverse of the PCr exponential recovery time constant, τ (kPCr= 1/τPCr). These values were then adjusted for PCr depletion.

### 6. Functional Assessment

Cardiorespiratory fitness was assessed by maximum oxygen consumption during peak exercise (peak VO_2_ or pVO_2_) during a modified treadmill Balke test (44). Gas exchange analyzer (MedGraphics, St. Paul, MN, USA) was used to capture the consumed O_2_ and expired CO_2_ concentrations, and oxygen consumption was calculated every 30 seconds. The highest oxygen consumption is the recorded peak VO_2_. Grip strength was measured using a hand-held dynamometer 3 times for each hand. The highest of each measure was used in the analysis.

### 7. Proteomics

Plasma proteomic profiles were assessed using SomaScan 1.3K as previously described (24,44–46). In brief, this is a multiplexed, aptamer (SOMAmer)-based assay that simultaneously measures thousands of human proteins broadly ranging from femto- to micro-molar concentrations. This technology relies on protein-capture SOMAmer (Slow Off-rate Modified Aptamer) reagents (47) designed to optimize high affinity, slow off-rate, and high specificity to target proteins. Proteins analyzed cover major molecular functions including receptors, kinases, growth factors, and hormones, and span a diverse collection of secreted, intracellular, and extracellular proteins or domains. The final data included relative abundances of 1305 SOMAmers or proteins expressed as relative fluorescence units (RFU). The raw data went through a series of normalization processes to account for hybridization, inter-sample, and inter-plate technical differences (24,45). Protein abundances were natural log-transformed, outliers outside 4SD were removed (0.47%) and were imputed using k-nearest neighbor (KNN).

### 8. Data processing and statistical analyses

### 8.1. Univariate statistical analysis

Correlation coefficients between respiratory parameters and age of GESTALT serum donor or epigenetic age of GESTALT serum donor (**Figures 1-2**) were determined using Pearson’s correlation. The association between individual SOMAmer abundance and bioenergetic effects of serum were computed using multivariable regression models (SOMAmer abundance = independent variable; bioenergetic effect = dependent variable) adjusted for sex and SomaScan batch (**Figure 3**, **Table 1**). A Bonferroni corrected P value of 3.83×10^−5^ (0.05/1305) was considered significant for the analysis of 1305 SOMAmers to adjust significance for multiple comparisons.

**FIGURE 1.**
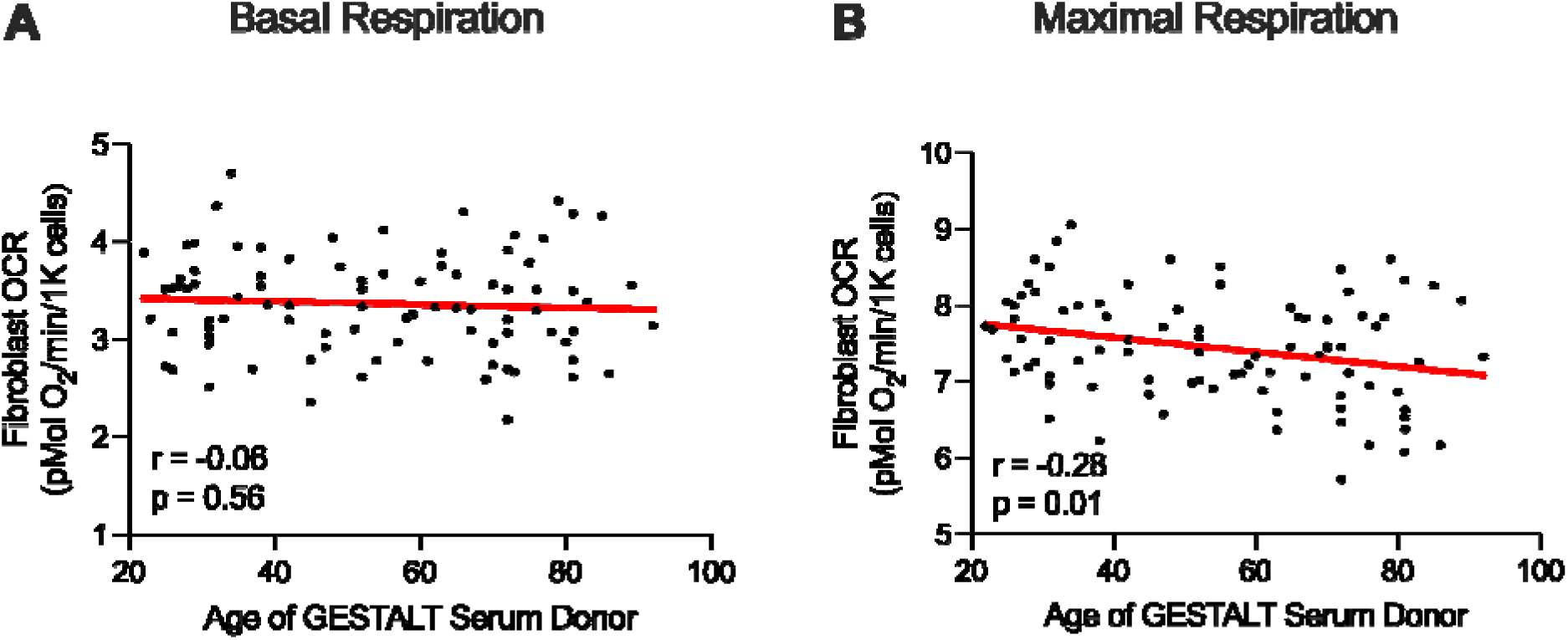
Maximal respiration, but not basal respiration, negatively correlates with the chronological age of the serum donor. N=88 GESTALT serum samples treated on human primary fibroblasts (fibroblast donor = male, 33 y.o).

**FIGURE 2.**
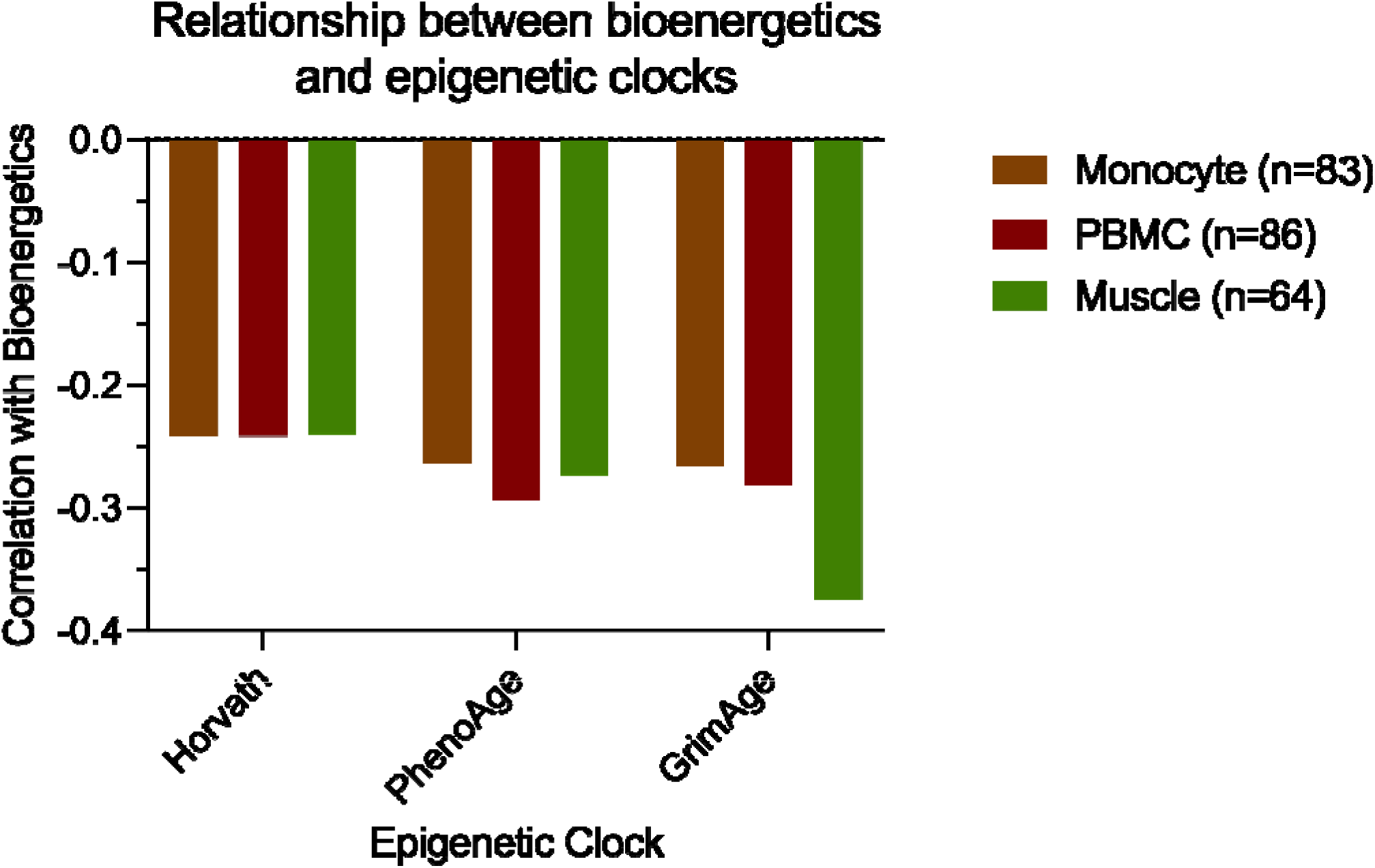
Serum-mediated bioenergetic effects relate to the epigenetic age of serum donors in multiple tissue types. Serum-mediated bioenergetic effects are based on the maximal respiration of young fibroblast donor.

**FIGURE 3.**
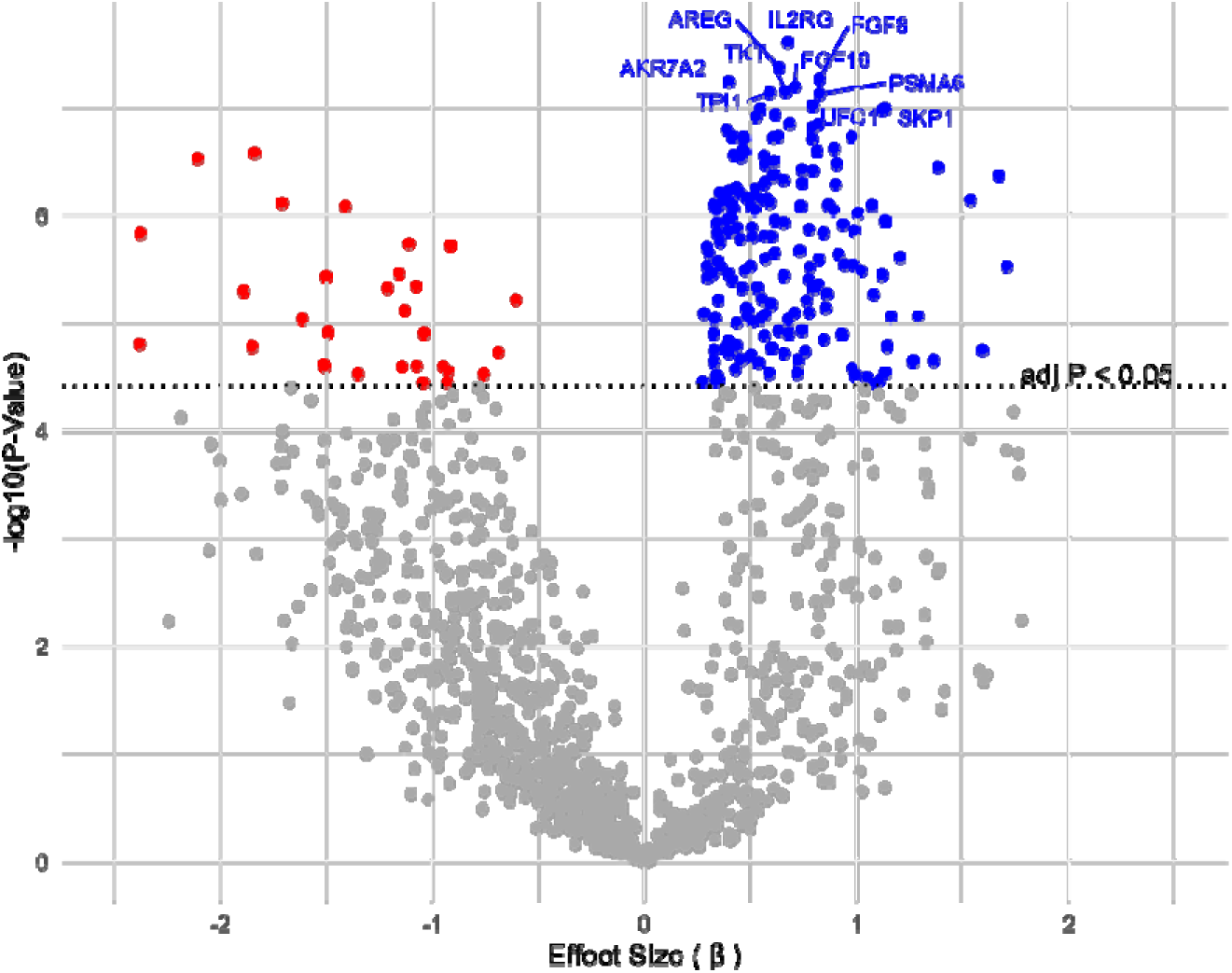
Serum proteins are positively (n=183; blue) and negatively (n=32; red) associated with the bioenergetic effects of serum using univariate linear regression models. Results are adjusted for sex and batch; used Bonferroni correction.

**TABLE 1:** Top 25 most significant proteins associated with maximal respiration of the young fibroblast donor treated with GESTALT serum. Protein targets (SOMAmers) are ranked by beta coefficients. Proteins in bold type indicate that they overlap with protein targets identified in the multivariate analysis.

| SeqId | Gene ID | UniProt | Target | Beta | SE | P value |
| --- | --- | --- | --- | --- | --- | --- |
| **2634-2** | **IL2RG** | **P31785** | **IL-2 sRg** | **0.6751** | **0.1093** | **2.47E-08** |
| **2970-60** | **AREG** | **P15514** | **AREG** | **0.6344** | **0.1048** | **4.23E-08** |
| **4394-71** | **FGF8** | **P55075** | **FGF-8A** | **0.8257** | **0.1378** | **5.43E-08** |
| 4188-1 | AKR7A2 | O43488 | Aflatoxin B1 aldehyde reductase | 0.3970 | 0.0664 | 5.76E-08 |
| **4393-3** | **FGF16** | **O43320** | **FGF-16** | **0.7094** | **0.1191** | **6.40E-08** |
| 4306-4 | TKT | P29401 | Transketolase | 0.6652 | 0.1121 | 7.07E-08 |
| 4309-59 | TPI1 | P60174 | Triosephosphate isomerase | 0.5899 | 0.0995 | 7.25E-08 |
| **3860-7** | **PSMA6** | **P60900** | **PSA6** | **0.8260** | **0.1395** | **7.46E-08** |
| **3405-6** | **UFC1** | **Q9Y3C8** | **UFC1** | **0.7931** | **0.1354** | **9.61E-08** |
| 3902-21 | SKP1 | P63208 | SKP1 | 1.1324 | 0.1938 | 1.02E-07 |
| **4963-19** | **ARPP19** | **P56211** | **ARP19** | **0.5466** | **0.0936** | **1.03E-07** |
| 4547-59 | FLRT1 | Q9NZU1 | FLRT1 | 1.1223 | 0.1923 | 1.06E-07 |
| 3897-61 | PDXP | Q96GD0 | PLPP | 0.6162 | 0.1060 | 1.16E-07 |
| **4249-64** | **NME2** | **P22392** | **NDP kinase B** | **0.5251** | **0.0905** | **1.22E-07** |
| **3874-8** | **UBE2L3** | **P68036** | **UB2L3** | **0.6844** | **0.1187** | **1.41E-07** |
| 5888-29 | EIF5A | P63241 | eIF-5A-1 | 0.8188 | 0.1420 | 1.43E-07 |
| **4389-2** | **DHH** | **O43323** | **DHH** | **0.7911** | **0.1376** | **1.53E-07** |
| 5346-24 | CPNE1 | Q99829 | CPNE1 | 0.3868 | 0.0674 | 1.60E-07 |
| **10363-13** | **SMAD3** | **P84022** | **SMAD3** | **0.6303** | **0.1105** | **1.85E-07** |
| 3593-72 | CASP3 | P42574 | Caspase-3 | 0.4143 | 0.0727 | 1.86E-07 |
| 4396-54 | IFNL1 | Q8IU54 | IFN-lambda 1 | 0.9770 | 0.1714 | 1.87E-07 |
| 3817-18 | PRKCQ | Q04759 | KPCT | 0.4678 | 0.0821 | 1.88E-07 |
| 4258-15 | PA2G4 | Q9UQ80 | PA2G4 | 0.6078 | 0.1067 | 1.90E-07 |
| 3877-67 | CAMKK1 | Q8N5S9 | CaMKK alpha | 0.7888 | 0.1386 | 1.94E-07 |
| 3848-14 | GAPDH | P04406 | GAPDH, liver | 0.4699 | 0.0827 | 2.01E-07 |

### 8.2. Construction of Proteomic Signature of Systemic Mitochondrial Health (MITOAGE)

We constructed a proteomic predictor of serum-mediated bioenergetics by fitting elastic net regression models using the eNetXplorer R package, version 1.1.3, publicly available at https://github.com/juliancandia/eNetXplorer (49). The elastic net mixing parameter α was tested from 0 (ridge) to 1 (lasso) in 0.1 intervals and utilized 500 five-fold cross-validation runs, each of them compared to 125 null model permutations. Within the elastic net, the cross-validations utilized an 80/20 split. Participant sex was included as a covariate. Significant features (*P* < 0.05) in the selected alpha model were identified as the proteomic signature (**Table 2**). Protein weights and predicted values were derived from a glmnet ridge regression. Using the selected proteins and weights, MitoAge was created as a linear combination of the products of the weight and the protein abundances. To evaluate the precision of MitoAge, we then examined the association between the predicted and actual serum-mediated bioenergetics. We further evaluated the association of MitoAge with serum-mediated bioenergetics in a separate fibroblast line as well as with an *in vivo* measures of mitochondrial OXPHOS (^31^P-MRS) with Pearson’s correlation. To evaluate how well MitoAge reflects additional age-related functional measures, we examined the association of MitoAge with pVO_2_ and grip strength, both functional measures known to decline with age. Our goal with MitoAge was to develop a signature of age-related mitochondrial health, therefore participant chronological age was not included as a covariate. To address how age may affect the model, we performed a causal mediation analysis to examine whether MitoAge mediated the association between age and bioenergetics. The mediator model regressed the MitoAge on age, and the outcome model regressed bioenergetics on both age and MitoAge. Indirect, direct, and total effects were estimated using nonparametric bootstrapping with 5,000 simulations and percentile-based 95% confidence intervals. Analyses were performed in R using the mediation package.

**TABLE 2:** MitoAge Features. Proteins (or sex) are listed in order of p-value (corresponding with Supplemental File 1). Proteins in bold type indicate that they overlap with protein targets identified in the univariate analysis.

| SeqId | GENE ID | UniProt | Target | Weight |
| --- | --- | --- | --- | --- |
| 2578-67 | CCL2 | P13500 | MCP-1 | -0.433 |
| **2970-60** | **AREG** | **P15514** | **AREG** | **-0.002** |
| 5508-62 | CTSD | P07339 | Cathepsin D | -0.45 |
| 3497-13 | IFNA2 | P01563 | IFN-aA | -0.443 |
| **2634-2** | **IL2RG** | **P31785** | **IL-2 sRg** | **0.025** |
| **10363-13** | **SMAD3** | **P84022** | **SMAD3** | **0.011** |
| 3232-28 | ACP5 | P13686 | TrATPase | -0.416 |
| 3059-50 | TNFSF13B | Q9Y275 | BAFF | -0.257 |
| **4393-3** | **FGF16** | **O43320** | **FGF-16** | **-0.009** |
| 3535-84 | DKK1 | O94907 | DKK1 | 0.164 |
| 2630-12 | IL1RAP | Q9NPH3 | IL-1 R AcP | 0.271 |
| 2826-53 | EDA | Q92838 | EDA | 0.275 |
| 5460-60 | DDX19B | Q9UMR2 | DEAD-box protein 19B | 0.052 |
| **4249-64** | **NME2** | **P22392** | **NDP kinase B** | **0.024** |
| 2944-66 | NBL1 | P41271 | DAN | -0.294 |
| 10344-334 | IL10RA | Q13651 | IL-10 Ra | -1.053 |
| **3860-7** | **PSMA6** | **P60900** | **PSA6** | **0.016** |
| 3810-50 | FGR | P09769 | FGR | -0.14 |
| **4389-2** | **DHH** | **O43323** | **DHH** | **0.036** |
| **3874-8** | **UBE2L3** | **P68036** | **UB2L3** | **0.084** |
| **3405-6** | **UFC1** | **Q9Y3C8** | **UFC1** | **0.107** |
| **4963-19** | **ARPP19** | **P56211** | **ARP19** | **0.006** |
| 4549-78 | FUT5 | Q11128 | FUT5 | -0.055 |
| **4394-71** | **FGF8** | **P55075** | **FGF-8A** | **0.118** |
| 5245-40 | PRKAA2 PRKAB2 PRKAG1 | P54646 O43741<br>P54619 | AMPK a2b2g1 | 0.01 |
|  |  |  | Intercept | 7.497 |
|  |  |  | Sex (Male) | -0.12 |

**TABLE 3:** Associations of bioenergetic and clinical parameters with MitoAge. The correlations between MitoAge and 1) serum-mediated maximal respiration of the old fibroblast donor, 2) ^31^P-MRS, 3) pVO_2_, and 4) grip strength are described. These are compared with the associations with serum-mediated maximal respiration of the young fibroblast donor. *kPCR via ^31^P-MRS is partialed for PCr depletion.

|  | MitoAge<br>unadjusted |  | Bioenergetics (33 y.o.<br>fibroblast donor) |  |
| --- | --- | --- | --- | --- |
|  | r | p | r | p |
| Bioenergetics (72 y.o.) | 0.676 | <.001 | 0.828 | <.001 |
| kPCR* | 0.251 | 0.024 | 0.244 | 0.028 |
| $\text{pVO}_2$ | 0.241 | 0.045 | 0.221 | 0.066 |
| Grip Strength | -0.008 | 0.944 | -0.048 | 0.667 |

## RESULTS

### Participant demographics

The participant ages in this analytical group were between 22-92 (mean: 54.14 years), with body mass index (BMI) ranging from 19.6-31.0 (mean 25.8), and 42% were female.

### Serum-mediated mitochondrial bioenergetics negatively correlate with the chronological ages of the serum donors

We measured the bioenergetic capacity of primary human fibroblasts (33 y.o. donor) treated with GESTALT serum samples to assess how extrinsic factors in serum may cause differences in bioenergetics of naïve cells. We observed no significant relationship between the chronological age of the GESTALT serum donor and fibroblast basal respiration after serum treatment (R = 0.06, *P* = 0.56; **Figure 1A**). We found a significant correlation between the age of the GESTALT serum donors and maximal respiration after serum treatment (R = −0.28, *P* = 0.01; **Figure 1B**).

### Serum-mediated bioenergetic effects relate to epigenetic age of serum donors in multiple tissue types

To investigate if the bioenergetic effects of serum were more strongly associated with biological age than chronological age, we compared the correlations between fibroblast serum-mediated bioenergetics and epigenetic ages derived from PBMCs, monocytes, and muscle tissue from GESTALT participants (**Figure 2**). Epigenetic ages were calculated for the Horvath (pan-tissue), DNAmPhenoAge, and GrimAge clocks, as these represent potentially different aspects of aging (molecular, phenotypic, or mortality-focused). We observed that the serum-mediated bioenergetic metric and the Horvath clock were negatively correlated at the same magnitude across all three cell types (R = −0.24). Second generation epigenetic clocks, DNAmPhenoAge and GrimAge, were more strongly negatively correlated to serum-mediated bioenergetics compared to the Horvath clock for all cell or tissue types. We observed similar correlations between serum-mediated bioenergetics and DNAmPhenoAge of monocytes, PBMCs, and muscle tissue (R = −0.26; R = −0.29; R = −0.27, respectively). With GrimAge, differences in the correlations with monocytes, PBMCs, and muscle tissue (R = −0.27; R = −0.28; R = −0.37, respectively) were notable. Overall, the correlations between bioenergetics and the Horvath clock and the PhenoAge clock were very consistent regardless of tissue type (i.e., between monocytes, PBMCs, and muscle tissue), whereas the correlations involving the GrimAge clock have more variation depending on the tissue from which methylation data were derived.

### Identification of circulating proteins related to serum-mediated bioenergetics

We used univariate linear regression models to evaluate the abundance of circulating proteins in plasma correlated with the serum-mediated bioenergetics. Of the 1305 proteins tested, 215 proteins were significantly associated with serum-mediated bioenergetics (Bonferroni corrected *P* < 3.83×10^−5^). There were 183 positively associated proteins and 32 negatively associated proteins, as depicted by the volcano plot in **Figure 3**. The top 25 proteins ranked by *P* value are presented in **Table 1**, with an expanded table of all 1305 proteins available in **Supplemental File 1**. The protein with the strongest association with serum-mediated bioenergetics was interleukin 2 receptor subunit gamma (IL-2 sRg, Beta = 0.6751, *P* < 0.001). In addition to IL-2 sRg, the top 10 most significant proteins were amphiregulin (AREG), fibroblast growth factor 8A (FGF-8A), Aflatoxin B1 aldehyde reductase (AKR7A2), fibroblast growth factor 16 (FGF-16), transketolase (TKT), triosephosphate isomerase (TPI1), proteasome subunit alpha type-6 (PSA6), ubiquitin-fold modifier conjugating enzyme 1 (UFC1), and S-phase kinase-associated protein 1 (SKP1).

### Age-related proteomic signature predicts the bioenergetic effects of circulating factors

To derive the MitoAge, we fit an elastic net regression model to identify a set of protein markers associated with the bioenergetic effect of serum on young fibroblasts. The elastic net model accuracy is depicted in **Supplemental Figure 1**, which shows the correlation between the out-of-bag prediction and actual maximal respiration for each participant. The selected elastic net mixing parameter alpha (0) and penalty parameter lambda (3.99) are also shown. The intercept is 7.497. The elastic net identified 25 proteins associated with maximal respiration (**Table 2**). The protein with the strongest association with serum-mediated bioenergetics was C-C motif chemokine 2 (MCP-1, Weight=-0.433, P=0.021). In addition to MCP-1, the top 10 most significant proteins identified from the elastic net included AREG, cathepsin D (CTSD), interferon alpha-2 (IFN-aA), IL-2 sRg, mothers against decapentaplegic homolog 3 (SMAD3), tartrate-resistant acid phosphatase type 5 (TrATPase), tumor necrosis factor ligand superfamily member 13B (BAFF), FGF-16, and Dickkopf-related protein 1 (DKK1).

The MitoAge final linear regression model parameters are displayed in **Supplemental File 1**. The correlation between the predicted values from the fitted MitoAge linear regression and actual serum-mediated bioenergetics was R = 0.84 (*P* < 0.001) (**Figure 4**). The mediation analysis indicated that MitoAge mediated the effect of age on bioenergetics, accounting for 91% of the total effect, demonstrating that the relationship between age and bioenergetics is largely explained by MitoAge (**Table 4**).

**FIGURE 4.**
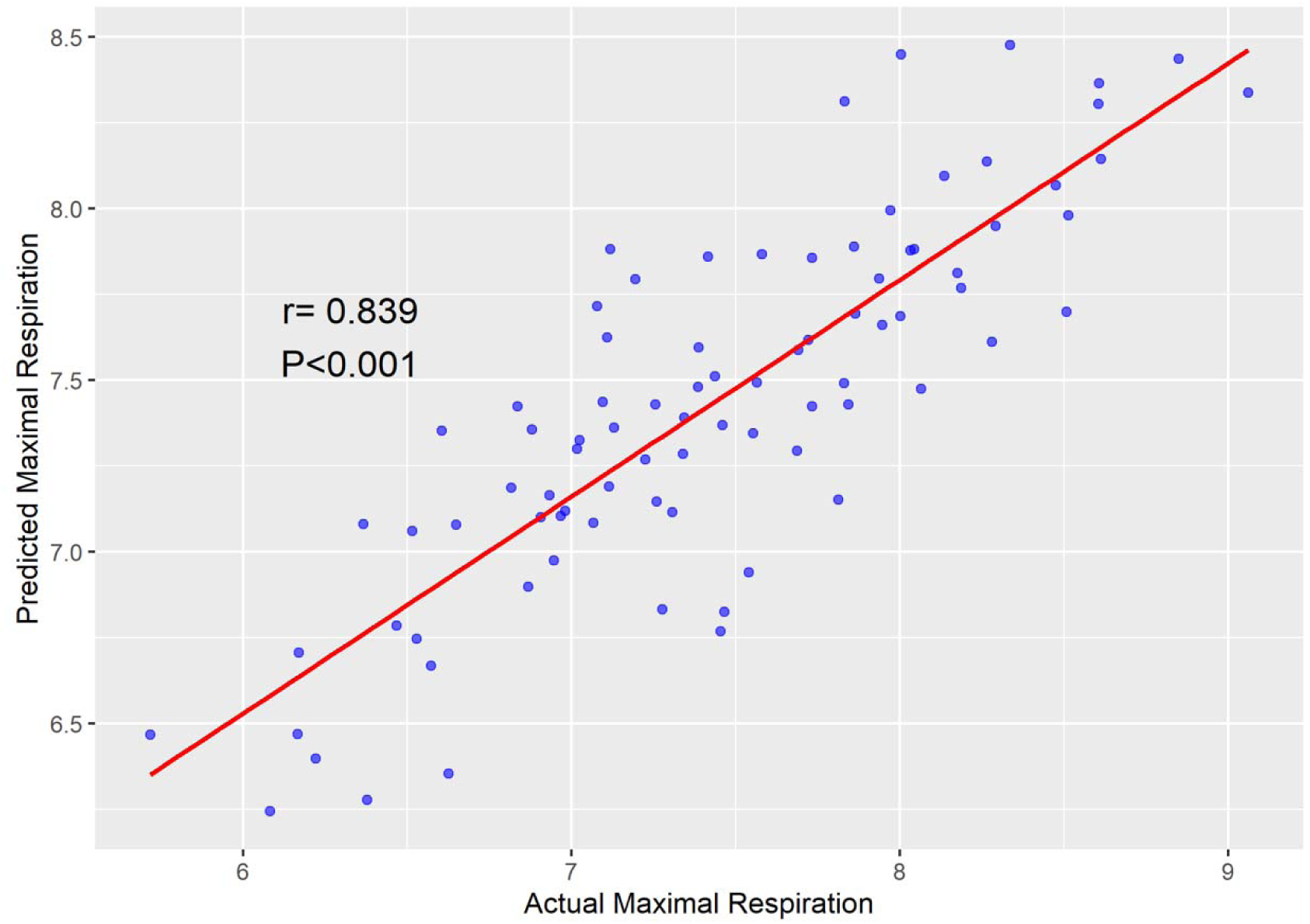
Precision and accuracy of MitoAge. Correlation between the actual (observed) maximal respiration from serum treatment versus the predicted maximal respiration using the proteomic signature.

**TABLE 4:** Mediation analysis to examine relationships between age and bioenergetics. Indirect, direct, and total effects are shown, and were estimated using nonparametric bootstrapping with 5,000 simulations and percentile-based 95% confidence intervals.

|  | Estimate | 95% CI | p-value |
| --- | --- | --- | --- |
| ACME (Average Causal Mediation Effect) | -0.009 | -0.016, -0.002 | 0.006 |
| ADE (Average Direct Effect) | -0.009 | -0.005, 0.003 | 0.697 |
| Total Effect | -0.010 | -0.018, -0.002 | 0.012 |
| Proportion Mediated | 0.913 | 0.513, 1.656 | 0.010 |

Out of these proteins identified by the elastic net, 11 of them overlapped with the univariate analysis (**Table 2**, bolded type). The overlapping proteins are: IL-2 sRg, UFC1, FGF-16, cAMP-regulated phosphoprotein 19 (ARP19), AREG, SMAD3, ubiquitin conjugating enzyme E2 L3 (UB2L3), desert hedgehog protein (DHH), PSA6, FGF-8A, and nucleoside diphosphate kinase B (NME2).

### Serum mediates bioenergetic differences regardless of fibroblast donor age

In order to assess the potential influence of intrinsic cellular age on serum-mediated bioenergetics, we repeated the serum exposure on fibroblasts from an older donor (72 y.o. donor, **Supplemental Figure 2)**. Similar to the serum treatment of young fibroblast cells (33 y.o. donor), we found a significant correlation between the age of the GESTALT serum donors and maximal respiration after serum treatment (R = −0.25, *P* = 0.02).

### Testing of MitoAge signature

We sought to validate MitoAge by using independent measures of bioenergetic capacity within the test sample. Therefore, we examined the association between MitoAge and the bioenergetic effect of the serum sample on fibroblasts derived from the older individual. We observed a positive correlation between the MitoAge and the actual observed respiration of the serum-treated fibroblasts from the older donor (R = 0.73, *P* < 0.001; **Figure 5A**; **Table 3**).

**FIGURE 5.**
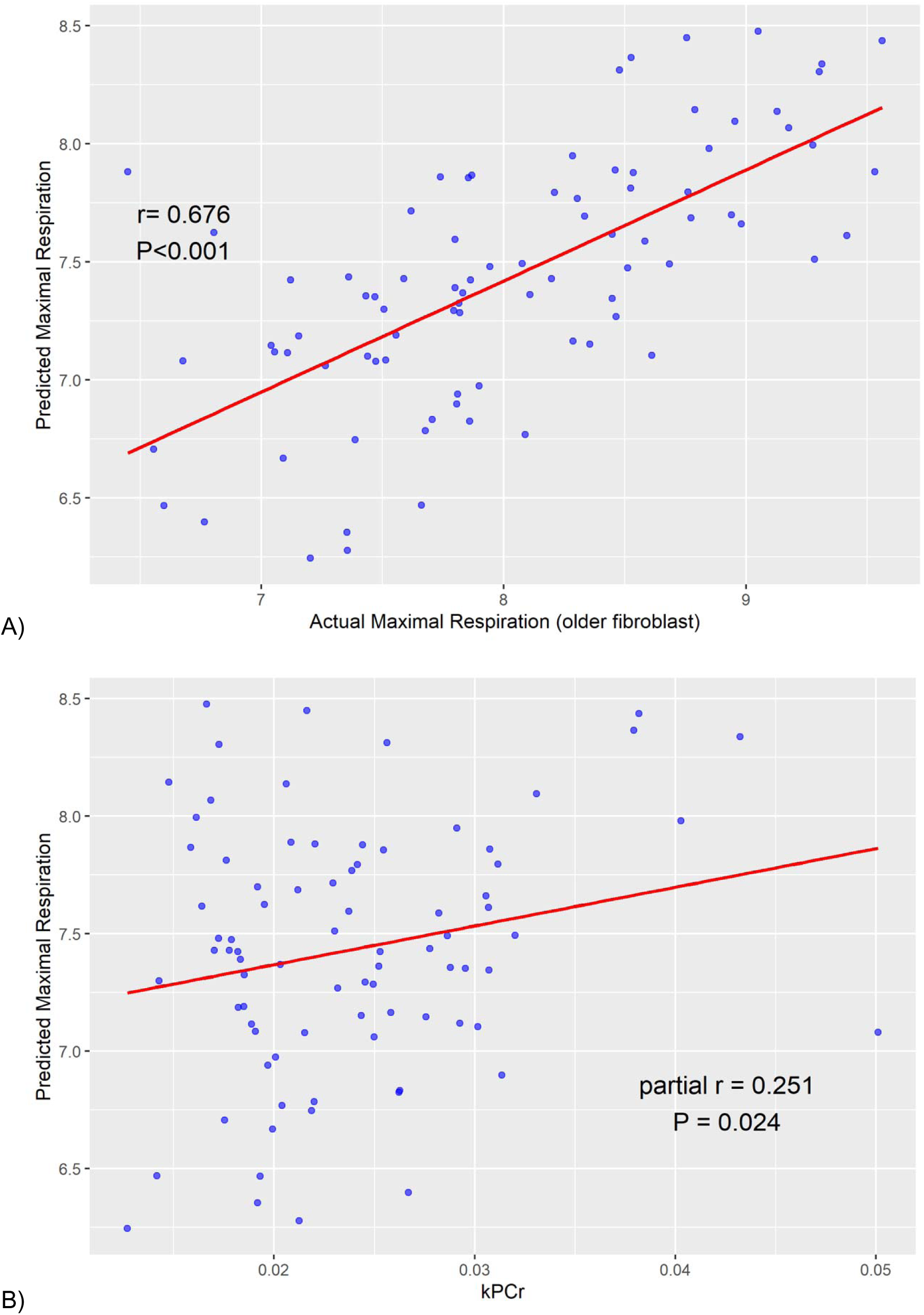
Validation of MitoAge. A) Correlation between the actual (observed) maximal respiration from serum treatment in older fibroblasts compared to the predicted maximal respiration of the young fibroblast donor using the proteomic signature. B) Correlation between the actual (observed) *in vivo* kPCr via ^31^P-MRS compared to the predicted maximal respiration of the young fibroblast donor using MitoAge.

### Prediction of *in vivo* mitochondrial function

We also used the measure of maximum oxidative capacity (quantified via ^31^P-MRS as post-exercise recovery rate of phosphocreatine “kPCr”; **Figure 5B**) to determine if MitoAge was associated with an *in vivo* measure of mitochondrial function. Our results indicated that MitoAge was positively correlated to kPCr (R = 0.25, *P* = 0.003; **Figure 5B**; **Table 3**).

### MitoAge is associated with cardiorespiratory fitness, but not strength

We evaluated correlations between MitoAge and measures of physical function, such as pVO_2_, and grip strength (**Table 3**). MitoAge had a positive correlation with pVO_2_ (R = 0.262, *P* = 0.028). We observed no correlation between MitoAge and grip strength (R = 0.022, *P* = 0.843).

## DISCUSSION

The results of this study demonstrate some compounds in serum can actively mediate age-related related bioenergetic differences in cultured fibroblasts regardless of the intrinsic age of the target cell. The negative relationship between the age of the GESTALT serum donor and fibroblast OCR post-serum treatment was similar regardless of fibroblast donor age. This is in line with the premise that differences in mitochondrial bioenergetics are primarily driven by non-cellular factors present in circulation. (15). It is notable that correlations between the bioenergetic effects of serum are more strongly related to the participants’ biological age, based on epigenetic clocks, rather than the chronological of the serum donor. This is supported by our finding that chronological age and the Horvath clock had similar correlations to the bioenergetic effects of serum. We believe this may be the case as first generation of epigenetic clocks (such as Horvath and Hannum) were trained to predict chronological age (38,50). Second generation clocks (such as DNAmPhenoAge and GrimAge) were more strongly associated with the bioenergetic effects of serum regardless of tissue type. These second generation clocks were designed to capture biological age by taking into account phenotypic differences and risks for mortality. DNAmPhenoAge was trained on markers of phenotypic age using clinical biomarkers, such as albumin, creatinine, and glucose (39), while GrimAge was trained on various protein biomarkers and mortality risk factors, such as years of smoking history – all of which affect biological age (40,51). These results add to a growing body of evidence that mitochondrial function may contribute to differences in biological aging.

It is also interesting to note the difference in strength of correlation between bioenergetics and epigenetic age among the tissues measured. The correlation between bioenergetics and the Horvath clock was similar among all tissues measured. This is likely due to the original, pan-tissue Horvath clock being trained on multiple tissues, therefore its age estimation is robust across multiple tissues. The other clocks were not necessarily designed to be pan-tissue, which is why there may be more subtle differences in age estimation across tissue types. In comparison, the correlation between bioenergetics and GrimAge was quite different among tissues. Of all correlations, GrimAge from muscle tissue had the strongest negative correlation with bioenergetics compared to that of PBMCs or monocytes. It is possible that this is the case because GrimAge includes protein surrogates, such as leptin and GDF-15, which may have greater age-related mitochondrial and metabolic relevance, especially in a more metabolically active tissues such as skeletal muscle.

Our univariate and multivariate analyses both identified proteins associated with the bioenergetic effects of the serum. Of the proteins that were common to both analyses, some have known functions in aging biology, mitochondrial function, and intercellular signaling, further supporting the idea that these circulating proteins can have broad, systemic effects on a wide range of tissues. AREG, for example, is a ligand of the EGF receptor and is associated with delayed cell death (52) and senescence in certain cell types (53). SMAD3 is an intracellular signal transducer and transcriptional modulator which was found to be reduced in older animals compared to young (54). Age-related changes in SMAD3 signaling can impact ROS production and has implications in neurodegeneration (54). Age-related SMAD3 signaling has also been associated with cell senescence (55) and pro-inflammatory factors (56).

While 11 proteins identified in this study were common to both univariate and multivariate analyses, many significant proteins were unique to each list. These discrepancies may arise from the different methods used in each approach: the multivariate analysis takes into account the simultaneous effects of multiple independent variables, thereby affecting the impact of individual explanatory variables on the response. It is important to recognize that most proteins were identified by one method, but not both. For example, MCP-1 was the protein identified by the elastic net to have the strongest association with serum-mediated bioenergetics, and was not identified by the univariate approach. MCP-1 modulates innate immune response and inflammatory defense and has also been shown in animal models to decrease lifespan and promote metabolic dysregulation (57,58). AKR7A2, identified by our univariate approach, has a strong association with bioenergetics, but was not identified by the elastic net. AKR7A2 also has metabolic effects associated with aging, such as in mediating oxidative stress (59,60). Both MCP-1 and AKR7A2 are implicated in age-related metabolic changes; therefore, our comprehensive approach using both univariate and multivariate methods allows us to identify proteins that may be missed by using only one method.

Of the 25 proteins identified in our signature, only two were previously identified in Tanaka et al.’s plasma proteomic signature of age, CTSD and ectodysplasin-A (EDA), which utilized proteomic data from the same study (24). It is reasonable that some proteins would be identified in both signatures. CTSD is associated with cognitive aging and neurodegeneration (61,62) and EDA is associated with pro-inflammatory phenotypes and muscle wasting associated with aging (63,64). The fact that only two out of the 25 proteins identified in our signature were previously identified in a chronological aging signature further corroborates the results of our mediation analysis showing that MitoAge mediates 91% of the effect of age on bioenergetics, with chronological age alone only accounting for 9% of this effect.

In this study, we demonstrate that circulating proteins can drive differences in mitochondrial function associated with age. This recognition enabled us to develop MitoAge as a powerful predictor of the bioenergetic effects of circulating factors that is also associated with complementary bioenergetic outcomes, including the recovery of ATP in skeletal muscle after exercise-induced depletion. Aligned with previous studies that have reported relationships between mitochondrial bioenergetic capacity with aging and physical performance (29,30,65), results presented here demonstrate that MitoAge is similarly associated with pVO_2_. Interestingly, MitoAge was not associated with grip strength, suggesting that strength may be most closely related to other factors associated with advancing age. Overall, MitoAge and the findings presented here align with previous literature indicating the strong interdependent relationship between mitochondrial function and physical fitness (30,66,67). Importantly, our analyses demonstrate that MitoAge can be utilized as a reliable biomarker of mitochondrial health in the context of human aging and functional decline.

This study is not without limitations. While this study utilizes cross-sectional serum samples from individuals of different ages, future studies may utilize longitudinal samples derived from individuals as they advance in age. Using MitoAge longitudinally could reflect the heterogeneity of aging trajectories both within and across individuals. Furthermore, larger studies applying MitoAge to larger cohorts are needed to investigate potential sex differences. Similar to other promising biomarkers of aging, the development of sex-specific markers and signatures should be evaluated.

Among the major innovations presented here is the use of a biologically-informed and hypothesis-driven computational approach to develop MitoAge. This study introduces a new proteomic signature built upon a widely recognized cellular hallmark of aging: mitochondrial dysfunction. Reliable markers that can inform on cellular hallmarks of aging represent a new generation of aging biomarkers. There are many known contributors to alterations in age-related mitochondrial OXPHOS, such as diet, exercise, fitness, environmental exposures, and cognitive ability (16,18,29,30,67,68). MitoAge can be used across multiple studies to inform on the role of age-related bioenergetic differences and broaden our understanding of drivers of health and longevity. Further, MitoAge can be used as an outcome in clinical trials of agents designed to promote healthy longevity by targeting mitochondrial dysfunction.

## ACKNOWLEDGEMENTS

We acknowledge the San Diego Nathan Shock center (SD-NSC) for access to primary human dermal fibroblasts. We also acknowledge Adrian Arciniega for his help organizing mitochondrial data. We are grateful to the GESTALT participants for their participation in this clinical study.

## COMPETING INTERESTS AND FUNDING SOURCES

This work was supported by the following grants awarded to Anthony Molina: R01 AG054523, R01 AG061805, R01 AG072734, P30 AG068635, and Wellcome Leap. Anthony Molina is employed by the University of California San Diego. This research was also supported in part by the Intramural Research Program of the National Institutes of Health (NIH). The contributions of the NIH authors were made as part of their official duties as NIH federal employees, are in compliance with agency policy requirements, and are considered Works of the United States Government. However, the findings and conclusions presented in this paper are those of the authors and do not necessarily reflect the views of the NIH or the U.S. Department of Health and Human Services.

## DATA AVAILABILITY STATEMENT

Data is provided within the manuscript or supplementary information files. Additional data files may be available upon request.

## ETHICS AND CONSENT STATEMENT

GESTALT study was approved by the institutional review board of the National Institutes of Health (NIH). Informed consent as well as the consent to publish the data collected was obtained from every participant. In addition, for the epigenetic clock data presented here, which is based on DNA methylation studies, all participants were required to consent to DNA/RNA testing and storage at all visits in order to participate in the study. The GESTALT IRB approval number is 15-AG-0063.

## AUTHOR CONTRIBUTIONS

SRH and AJAM conceptualized and designed the study. SRH and NSS performed biological assays. SRH and JNB performed statistical analyses. JNB performed computational approaches in machine learning. QT, AZM, TT and MK assisted in collecting and organizing clinical data. BC and JC provided computational and methodological support with multi-omic data. AB, CD, GF, and MK assisted with data acquisition and blood isolation. RS and LF conceptualized and designed the GESTALT study. SRH wrote the first draft of the manuscript, which was edited and revised by SRH, TT, JC, AJAM, and LF. All authors read and approved the submitted version.

## SUPPLEMENTAL FIGURES AND TABLES

**Supplemental Figure 1:**
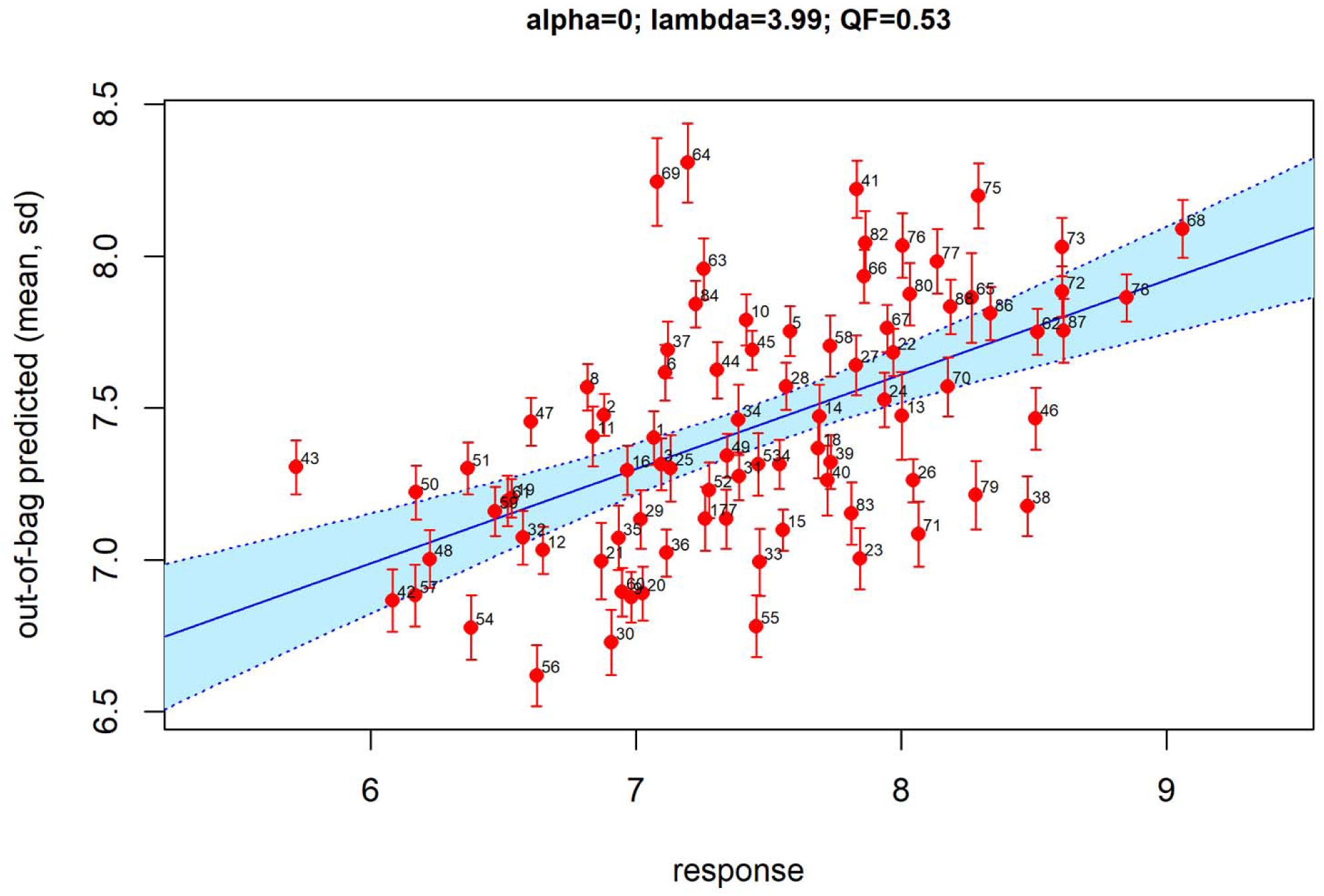
Computational methods for elastic net model accuracy. An elastic net model was assessed using eNetXplorer (1) with parameters n_run=500, n_perm_null=125, n_fold=5. Using Pearson’s correlation between the actual maximal respiration (x-axis) and out-of-bag predicted rmaximal respiration (y-axis) as quality factor (QF), the optimal model was defined by mixing parameter alpha=0 and shrinkage parameter lambda=3.99. Symbols represent individual participants; error bars represent the range of variation (standard deviation) of predicted values.

---

**Supplemental Figure 2:**
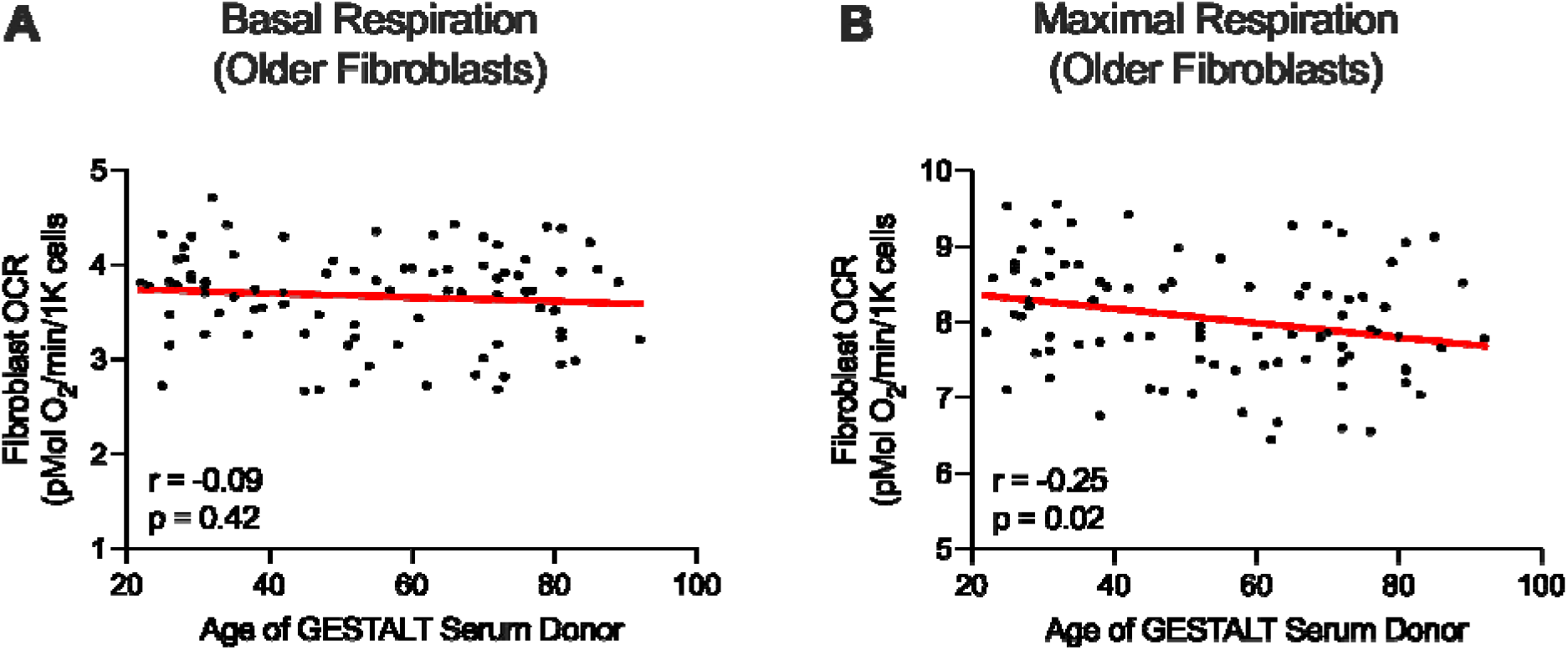
Maximal respiration, but not basal respiration, negatively correlates with the chronological age of the serum donor. N=88 GESTALT serum samples treated on human primary fibroblasts (fibroblast donor = male, 72 y.o).

## REFERENCES

1. López-Otín C, Blasco MA, Partridge L, Serrano M, Kroemer G. Hallmarks of aging: An expanding universe. Cell. 2023 Jan;186(2):243–78.

2. Serviddio G, Bellanti F, Romano AD, Tamborra R, Rollo T, Altomare E, et al. Bioenergetics in aging: mitochondrial proton leak in aging rat liver, kidney and heart. Redox Rep. 2007 Feb;12(1–2):91–5.

3. Kujoth GC, Hiona A, Pugh TD, Someya S, Panzer K, Wohlgemuth SE, et al. Mitochondrial DNA Mutations, Oxidative Stress, and Apoptosis in Mammalian Aging. Science. 2005 Jul 15;309(5733):481–4.

4. Carey BW, Finley LWS, Cross JR, Allis CD, Thompson CB. Intracellular α-ketoglutarate maintains the pluripotency of embryonic stem cells. Nature. 2015 Feb;518(7539):413–6.

5. Kaplon J, Zheng L, Meissl K, Chaneton B, Selivanov VA, Mackay G, et al. A key role for mitochondrial gatekeeper pyruvate dehydrogenase in oncogene-induced senescence. Nature. 2013 Jun;498(7452):109–12.

6. Tian Q, Mitchell BA, Zampino M, Fishbein KW, Spencer RG, Ferrucci L. Muscle mitochondrial energetics predicts mobility decline in well-functioning older adults: The baltimore longitudinal study of aging. Aging Cell. 2022 Feb;21(2):e13552.

7. Tian Q, Bilgel M, Walker KA, Moghekar AR, Fishbein KW, Spencer RG, et al. Skeletal muscle mitochondrial function predicts cognitive impairment and is associated with biomarkers of Alzheimer’s disease and neurodegeneration. Alzheimers Dement J Alzheimers Assoc. 2023 Oct;19(10):4436–45.

8. Tyrrell DJ, Bharadwaj MS, Jorgensen MJ, Register TC, Molina AJA. Blood cell respirometry is associated with skeletal and cardiac muscle bioenergetics: Implications for a minimally invasive biomarker of mitochondrial health. Redox Biol. 2016 Dec;10:65–77.

9. Tyrrell DJ, Bharadwaj MS, Jorgensen MJ, Register TC, Shively C, Andrews RN, et al. Blood-Based Bioenergetic Profiling Reflects Differences in Brain Bioenergetics and Metabolism. Oxid Med Cell Longev. 2017;2017:7317251.

10. Braganza A, Corey CG, Santanasto AJ, Distefano G, Coen PM, Glynn NW, et al. Platelet bioenergetics correlate with muscle energetics and are altered in older adults. JCI Insight. 2019 Jul 11;4(13):e128248.

11. Brack AS, Conboy MJ, Roy S, Lee M, Kuo CJ, Keller C, et al. Increased Wnt Signaling During Aging Alters Muscle Stem Cell Fate and Increases Fibrosis. Science. 2007 Aug 10;317(5839):807–10.

12. Conboy IM, Conboy MJ, Wagers AJ, Girma ER, Weissman IL, Rando TA. Rejuvenation of aged progenitor cells by exposure to a young systemic environment. Nature. 2005 Feb;433(7027):760–4.

13. Loffredo FS, Steinhauser ML, Jay SM, Gannon J, Pancoast JR, Yalamanchi P, et al. Growth Differentiation Factor 11 Is a Circulating Factor that Reverses Age-Related Cardiac Hypertrophy. Cell. 2013 May;153(4):828–39.

14. Salpeter SJ, Khalaileh A, Weinberg-Corem N, Ziv O, Glaser B, Dor Y. Systemic Regulation of the Age-Related Decline of Pancreatic β-Cell Replication. Diabetes. 2013 Aug 1;62(8):2843–8.

15. Gonzalez-Armenta JL, Li N, Lee RL, Lu B, Molina AJA. Heterochronic Parabiosis: Old Blood Induces Changes in Mitochondrial Structure and Function of Young Mice. J Gerontol A Biol Sci Med Sci. 2021 Feb 25;76(3):434–9.

16. Gonzalez-Armenta JL, Bergstrom J, Lee J, Furdui CM, Nicklas BJ, Molina AJA. Serum factors mediate changes in mitochondrial bioenergetics associated with diet and exercise interventions. GeroScience. 2023 Jun 27;46(1):349–65.

17. Amick KA, Mahapatra G, Gao Z, Dewitt A, Craft S, Jain M, et al. Plasma glycocholic acid and linoleic acid identified as potential mediators of mitochondrial bioenergetics in Alzheimer’s dementia. Front Aging Neurosci. 2022;14:954090.

18. Heimler SR, Amick KA, Bergstrom J, Moliné M, Mahapatra G, Craft S, et al. Nervonic acid and 15-epi-PGA1 mediate systemic mitochondrial dysfunction in AD dementia. GeroScience [Internet]. 2025 Jul 11 [cited 2025 Jul 16]; Available from: 10.1007/s11357-025-01776-6

19. Kawanishi N, Kato Y, Yokozeki K, Sawada S, Sakurai R, Fujiwara Y, et al. Effects of aging on serum levels of lipid molecular species as determined by lipidomics analysis in Japanese men and women. Lipids Health Dis. 2018 Jun 6;17(1):135.

20. Montoliu I, Scherer M, Beguelin F, DaSilva L, Mari D, Salvioli S, et al. Serum profiling of healthy aging identifies phospho- and sphingolipid species as markers of human longevity. Aging. 2014 Jan 21;6(1):9–25.

21. Yu Z, Zhai G, Singmann P, He Y, Xu T, Prehn C, et al. Human serum metabolic profiles are age dependent. Aging Cell. 2012;11(6):960–7.

22. Menni C, Kiddle SJ, Mangino M, Viñuela A, Psatha M, Steves C, et al. Circulating Proteomic Signatures of Chronological Age. J Gerontol A Biol Sci Med Sci. 2015 Jul;70(7):809–16.

23. Sebastiani P, Federico A, Morris M, Gurinovich A, Tanaka T, Chandler KB, et al. Protein signatures of centenarians and their offspring suggest centenarians age slower than other humans. [cited 2025 Mar 5]; Available from: https://onlinelibrary.wiley.com/doi/10.1111/acel.13290

24. Tanaka T, Biancotto A, Moaddel R, Moore AZ, Gonzalez-Freire M, Aon MA, et al. Plasma proteomic signature of age in healthy humans. Aging Cell. 2018 Oct;17(5):e12799.

25. Sun BB, Maranville JC, Peters JE, Stacey D, Staley JR, Blackshaw J, et al. Genomic atlas of the human plasma proteome. Nature. 2018 Jun;558(7708):73–9.

26. Tanaka T, Basisty N, Fantoni G, Candia J, Moore AZ, Biancotto A, et al. Plasma proteomic biomarker signature of age predicts health and life span. eLife. 2020 Nov 19;9:e61073.

27. Baird AL, Westwood S, Lovestone S. Blood-Based Proteomic Biomarkers of Alzheimer’s Disease Pathology. Front Neurol. 2015 Nov 16;6:236.

28. Di Narzo AF, Telesco SE, Brodmerkel C, Argmann C, Peters LA, Li K, et al. High-Throughput Characterization of Blood Serum Proteomics of IBD Patients with Respect to Aging and Genetic Factors. PLoS Genet. 2017 Jan 27;13(1):e1006565.

29. Tyrrell DJ, Bharadwaj MS, Van Horn CG, Marsh AP, Nicklas BJ, Molina AJA. Blood-cell bioenergetics are associated with physical function and inflammation in overweight/obese older adults. Exp Gerontol. 2015 Oct;70:84–91.

30. Tyrrell DJ, Bharadwaj MS, Van Horn CG, Kritchevsky SB, Nicklas BJ, Molina AJA. Respirometric Profiling of Muscle Mitochondria and Blood Cells Are Associated With Differences in Gait Speed Among Community-Dwelling Older Adults. J Gerontol A Biol Sci Med Sci. 2015 Nov;70(11):1394–9.

31. Kramer PA, Coen PM, Cawthon PM, Distefano G, Cummings SR, Goodpaster BH, et al. Skeletal Muscle Energetics Explain the Sex Disparity in Mobility Impairment in the Study of Muscle, Mobility and Aging. J Gerontol Ser A. 2024 Apr 1;79(4):glad283.

32. Shadel GS, Adams PD, Berggren WT, Diedrich JK, Diffenderfer KE, Gage FH, et al. The San Diego Nathan Shock Center: tackling the heterogeneity of aging. GeroScience. 2021 Oct;43(5):2139–48.

33. Phang HJ, Heimler SR, Scandalis LM, Wing D, Moran R, Nichols JF, et al. Protocol for the San Diego Nathan Shock Center Clinical Cohort: a new resource for studies of human aging. BMJ Open. 2024 Jun;14(6):e082659.

34. Roy R, Kuo PL, Candia J, Sarantopoulou D, Ubaida-Mohien C, Hernandez D, et al. Epigenetic signature of human immune aging in the GESTALT study. Belz GT, Isales C, editors. eLife. 2023 Aug 17;12:e86136.

35. Ubaida-Mohien C, Lyashkov A, Gonzalez-Freire M, Tharakan R, Shardell M, Moaddel R, et al. Discovery proteomics in aging human skeletal muscle finds change in spliceosome, immunity, proteostasis and mitochondria. Kaeberlein M, Rosen CJ, Kaeberlein M, Anderson R, editors. eLife. 2019 Oct 23;8:e49874.

36. Zhou W, Triche TJ Jr, Laird PW, Shen H. SeSAMe: reducing artifactual detection of DNA methylation by Infinium BeadChips in genomic deletions. Nucleic Acids Res. 2018 Nov 16;46(20):e123.

37. Zhou W, Laird PW, Shen H. Comprehensive characterization, annotation and innovative use of Infinium DNA methylation BeadChip probes. Nucleic Acids Res. 2017 Feb 28;45(4):e22.

38. Horvath S. DNA methylation age of human tissues and cell types. Genome Biol. 2013;14(10):R115.

39. Levine ME, Lu AT, Quach A, Chen BH, Assimes TL, Bandinelli S, et al. An epigenetic biomarker of aging for lifespan and healthspan. Aging. 2018 Apr 18;10(4):573–91.

40. Lu AT, Quach A, Wilson JG, Reiner AP, Aviv A, Raj K, et al. DNA methylation GrimAge strongly predicts lifespan and healthspan. Aging. 2019 Jan 21;11(2):303–27.

41. Choi S, Reiter DA, Shardell M, Simonsick EM, Studenski S, Spencer RG, et al. 31P Magnetic Resonance Spectroscopy Assessment of Muscle Bioenergetics as a Predictor of Gait Speed in the Baltimore Longitudinal Study of Aging. [cited 2025 Mar 11]; Available from: 10.1093/gerona/glw059

42. Zane AC, Reiter DA, Shardell M, Cameron D, Simonsick EM, Fishbein KW, et al. Muscle strength mediates the relationship between mitochondrial energetics and walking performance. Aging Cell. 2017;16(3):461–8.

43. Naressi A, Couturier C, Castang I, de Beer R, Graveron-Demilly D. Java-based graphical user interface for MRUI, a software package for quantitation of in vivo/medical magnetic resonance spectroscopy signals. Comput Biol Med. 2001 Jul 1;31(4):269–86.

44. Gold L, Ayers D, Bertino J, Bock C, Bock A, Brody EN, et al. Aptamer-Based Multiplexed Proteomic Technology for Biomarker Discovery. PLOS ONE. 2010 Dec 7;5(12):e15004.

45. Candia J, Daya GN, Tanaka T, Ferrucci L, Walker KA. Assessment of variability in the plasma 7k SomaScan proteomics assay. Sci Rep. 2022 Oct 13;12(1):17147.

46. Candia J, Cheung F, Kotliarov Y, Fantoni G, Sellers B, Griesman T, et al. Assessment of Variability in the SOMAscan Assay. Sci Rep. 2017 Oct 27;7(1):14248.

47. Rohloff JC, Gelinas AD, Jarvis TC, Ochsner UA, Schneider DJ, Gold L, et al. Nucleic Acid Ligands With Protein-like Side Chains: Modified Aptamers and Their Use as Diagnostic and Therapeutic Agents. Mol Ther Nucleic Acids. 2014 Oct 7;3(10):e201.

49. Candia J, Tsang JS. eNetXplorer: an R package for the quantitative exploration of elastic net families for generalized linear models. BMC Bioinformatics. 2019 Apr 16;20(1):189.

50. Hannum G, Guinney J, Zhao L, Zhang L, Hughes G, Sadda S, et al. Genome-wide Methylation Profiles Reveal Quantitative Views of Human Aging Rates. Mol Cell. 2013 Jan;49(2):359–67.

51. McCrory C, Fiorito G, Hernandez B, Polidoro S, O’Halloran AM, Hever A, et al. GrimAge Outperforms Other Epigenetic Clocks in the Prediction of Age-Related Clinical Phenotypes and All-Cause Mortality. J Gerontol Ser A. 2021 May 1;76(5):741–9.

52. Xu Q, Long Q, Zhu D, Fu D, Zhang B, Han L, et al. Targeting amphiregulin (AREG) derived from senescent stromal cells diminishes cancer resistance and averts programmed cell death 1 ligand (PD-L1)-mediated immunosuppression. Aging Cell. 2019;18(6):e13027.

53. Pommer M, Kuphal S, Bosserhoff AK. Amphiregulin Regulates Melanocytic Senescence. Cells. 2021 Feb;10(2):326.

54. Tichauer JE, Flores B, Soler B, Eugenín-von Bernhardi L, Ramírez G, von Bernhardi R. Age-dependent changes on TGFβ1 Smad3 pathway modify the pattern of microglial cell activation. Brain Behav Immun. 2014 Mar 1;37:187–96.

55. Gao XY, Lai YY, Luo XS, Peng DW, Li QQ, Zhou HS, et al. Acetyltransferase p300 regulates atrial fibroblast senescence and age-related atrial fibrosis through p53/Smad3 axis. Aging Cell. 2023;22(1):e13743.

56. Liu Y, Yu L, Xu Y, Tang X, Wang X. Substantia nigra Smad3 signaling deficiency: relevance to aging and Parkinson’s disease and roles of microglia, proinflammatory factors, and MAPK. J Neuroinflammation. 2020 Nov 16;17(1):342.

57. Luciano-Mateo F, Cabré N, Baiges-Gaya G, Fernández-Arroyo S, Hernández-Aguilera A, Elisabet Rodríguez-Tomàs E, et al. Systemic overexpression of C-C motif chemokine ligand 2 promotes metabolic dysregulation and premature death in mice with accelerated aging. Aging. 2020 Oct 26;12(20):20001–23.

58. Guo D, Zhu W, Qiu H. C-C Motif Chemokine Ligand 2 and Chemokine Receptor 2 in Cardiovascular and Neural Aging and Aging-Related Diseases. Int J Mol Sci. 2024 Jan;25(16):8794.

59. Ahmed MME, Wang T, Luo Y, Ye S, Wu Q, Guo Z, et al. Aldo-keto reductase-7A protects liver cells and tissues from acetaminophen-induced oxidative stress and hepatotoxicity. Hepatology. 2011;54(4):1322–32.

60. Wang Q, Lu T, Song P, Dong Y, Dai C, Zhang W, et al. Glycyrrhizic acid ameliorates hepatic fibrosis by inhibiting oxidative stress via AKR7A2. Phytomedicine. 2024 Oct 1;133:155878.

61. Vidoni C, Follo C, Savino M, Melone MAB, Isidoro C. The Role of Cathepsin D in the Pathogenesis of Human Neurodegenerative Disorders. Med Res Rev. 2016;36(5):845–70.

62. Urbanelli L, Emiliani C, Massini C, Persichetti E, Orlacchio A, Pelicci G, et al. Cathepsin D expression is decreased in Alzheimer’s disease fibroblasts. Neurobiol Aging. 2008 Jan 1;29(1):12–22.

63. Barbera MC, Guarrera L, Re Cecconi AD, Cassanmagnago GA, Vallerga A, Lunardi M, et al. Increased ectodysplasin-A2-receptor EDA2R is a ubiquitous hallmark of aging and mediates parainflammatory responses. Nat Commun. 2025 Feb 23;16(1):1898.

64. Özen SD, Kir S. Ectodysplasin A2 receptor signaling in skeletal muscle pathophysiology. Trends Mol Med. 2024 May 1;30(5):471–83.

65. Heimler S, Phang H, Bergstrom J, Mahapatra G, Dozier, Stephen, Gnaiger E, et al. Platelet bioenergetics are associated with resting metabolic rate and exercise capacity in older adult women. 2022 [cited 2023 Jan 4]; Available from: https://www.bioenergetics-communications.org/index.php/bec/article/view/Heimler_2022

66. Effects of physical activity and age on mitochondrial function | QJM: An International Journal of Medicine | Oxford Academic [Internet]. [cited 2025 Apr 30]. Available from: https://academic.oup.com/qjmed/article-abstract/89/4/251/1499967

67. Grevendonk L, Connell NJ, McCrum C, Fealy CE, Bilet L, Bruls YMH, et al. Impact of aging and exercise on skeletal muscle mitochondrial capacity, energy metabolism, and physical function. Nat Commun. 2021 Aug 6;12(1):4773.

68. Reddam A, McLarnan S, Kupsco A. Environmental Chemical Exposures and Mitochondrial Dysfunction: a Review of Recent Literature. Curr Environ Health Rep. 2022 Dec 1;9(4):631–49.

## References

1. Candia J, Tsang JS. eNetXplorer: an R package for the quantitative exploration of elastic net families for generalized linear models. BMC Bioinformatics. 2019 Apr 16;20(1):189.

